# Deciphering therapeutic efficacy of mycosynthesized silver nanoparticles using entomopathogenic fungi *Metarhizium anisopliae* against MCF-7 breast cancer cells *in vitro*

**DOI:** 10.1101/2022.08.25.505221

**Authors:** Sujisha S Nambiar, Arman Mohanty, Arupam Patra, Gurvinder Kaur Saini

## Abstract

Green synthesis of silver nanoparticles has gained much interest over few decades in the field of biomedical research due to their ease of synthesis, cost effectiveness, enhanced bioactivity and biocompatibility compared to the chemical synthesis. Recent studies on silver nanoparticles have shown their potential in various fields like antimicrobial, anticancer, larvicidal, catalytic, and wound healing properties. In the present study, entomopathogenic fungi *Metarhizium anisopliae* was used to synthesize silver nanoparticles. These silver nanoparticles were synthesized and characterized using UV-Vis spectroscopy, FESEM, FETEM and FTIR. Compared to the chemically synthesized silver nanoparticles, the mycosynthesized silver nanoparticles (MaAgNP) showed high yield. The size of mycosynthesized silver nanoparticles was found to be 5-20 nm and was spherical in shape. FTIR results confirmed the possible functional groups that are responsible for the reduction of silver ions. The mycosynthesized silver nanoparticles showed cytotoxicity on human breast cancer cell line (MCF-7) and the calculated IC_50_ value for MaAgNP was found to be 16.50 μg ml^-1^ whereas the chemically synthesized silver nanoparticles showed cytotoxicity but with increasing concentration, no further significant reduction in cell viability was observed. The possible reason behind improved cytotoxicity of MaAgNP can be the presence of extracellular secondary metabolites present in the fungal filtrate used to synthesize the nanoparticles. The MaAgNP was also observed to induce cell death through reactive oxygen species (ROS) generation.

## Introduction

Kingdom fungi are eukaryotic organisms which contain 2.2 million to 3.8 million species including microorganisms such as yeasts, molds, rusts, mildews, and mushrooms (David L Hawksworth & Robert Lucking, n.d.). The majority of fungus can be found living in soil or water, while others have either a parasitic or symbiotic association with other organisms, most commonly plants or animals. Mycelia are the bodies of fungi, which grow from the tips of filaments called hyphae. Fungi consume organic substances and absorb it into their mycelia (Feofilova, 2001). Fungi reproduce asexually or sexually by forming spores. The fungi that are capable to cause disease in arthropods are called entomopathogenic fungi. They usually enter the insect body in the form of spores (also called conidia) by penetrating the insect exoskeleton (Verena seidl, 2008). The entomopathogenicity of fungus can be utilized for the biocontrol of pests and insects that are harmful to agricultural crops. Myco pesticides are environment friendly and thus can substitute harmful chemical pesticides which cause challenges in soil–water purity and food safety. Spraying of fungal spore suspension is sufficient to cause infection in insect hosts and does not require ingestion which makes them a more attractive pest control agent. Fungi like *Beauveria bassiana, Beauveria brongniartii, Metarhizium anisopliae, Metarhizium rileyi and Isaria fumosorosea* are known to be entomopathogenic and do not cause infection in plants and animals. These fungi produce various secondary metabolites which are cytotoxic and some of them also have anti-cancerous properties. The organic compounds present in the secondary metabolites of fungi as well as the fungal enzymes has the ability to reduce the metal ions present in its salt solution to synthesize metallic nanoparticles (Chandra et al., 2013).

Nanotechnology is an evolving field due to its wide applications in various fields including therapeutics, pharmacology, biology, material science, physics, and chemistry. The conventional methods that are used to synthesize nanoparticles have disadvantages like low yield, high energy consumption, usage of hazardous chemicals, toxic by-products, high cost, etc (Abou El-Nour et al., 2010). To overcome these problems, green synthesis methods are utilized to synthesise nanoparticles. This green synthesis or biological synthesis of nanoparticles is done by using bacteria, fungi, actinomycetes, algae, plants and plant extracts. Among these methods, fungal-based synthesis of nanoparticles needs more attention, because of their higher productivity, easier handling, and low cost(Sastry et al., 2003.).

Silver is a noble metal, and in recent years, researchers have paid increasing attention to silver nanoparticles because of their unique qualities, such as their high electrical conductivity, chemical stability, catalytic activity, and antibacterial effects.. Recent studies show that mycosynthesised silver nanoparticles have anticancerous and insecticidal properties which can be utilised for cancer therapy and drug development (Shameli et al., 2010).

In this study, we have shown green synthesis of silver nanoparticles using the fungal extract of entomopathogenic fungi *Metarhizium anisopliae* which has improved cytotoxic potential through generation of reactive oxygen species (ROS) as compared to the chemically synthesized one.

## Materials and methods

### Fungal growth and maintenance

*Metarizium anisopliae* (MTCC 892) cultures were obtained from Microbial Type Culture Collection and Gene Bank (MTCC, India) and maintained on Potato dextrose agar medium at 28 °C for 10 days until sporulation.

### Chemical synthesis of AgNP

The silver nanoparticles were prepared by using a chemical reduction method (Gudikandula & Charya Maringanti, 2016). All reactants were prepared in double-distilled water. In a typical experiment, 100 ml of 1mM AgNO3 is boiled. In this solution, 5 ml of 1% trisodium citrate was added drop by drop. The solution was mixed vigorously during this process and heated until noticeable colour change (pale brown). The mixture was then taken from the heating element and swirled until room temperature.

### Mycosynthesis of silver nanoparticles

Fungal spores were collected by using a 0.01% Triton-X 100 solution. 10μl spore suspension (1*10^6^ spores/ml) was inoculated in 100ml Potato dextrose broth and incubated at 28°C for 72 hours. The fungal biomass was collected by centrifugation (5000 rpm, 10min), then washed two times with sterile double distilled water followed by centrifugation to remove all the media components. Fungal biomass (~10g) was inoculated in 100ml double distilled water and was incubated at 28 °C for 72 hours in an orbital shaker (120 rpm). After the incubation, the fungal filtrate was collected by centrifugation (5000 rpm, 10min). To the fungal filtrate, 16.9 mg of Silver nitrate (AgNO3) was added to make 1 mM AgNO_3_ solution and incubated at 28°C and 120 rpm in an orbital shaker in dark conditions until the brownish-yellow colour formation was observed. Control was maintained without the addition of silver nitrate (AgNO3) in the fungal filtrate (Amerasan et al., 2016).

### Characterization of bioactive nanoparticles

#### UV-VIS Spectroscopy

Exploiting the active interaction of silver nanoparticles with specific wavelengths of light, both biologically and chemically synthesized silver nanoparticles were monitored using Thermo scientific evolution 220 UV–VIS spectrophotometer (Zhang et al., 2016).

#### Field Emission Scanning Electron Microscope (FESEM)

Utilizing a scanning electron microscope, the nanoparticle’s size, shape, and morphologies were determined. FESEM provides high-resolution pictures of the sample’s surface. The sample was analyzed using FESEM, Zeiss, and Model: Sigma 300 microscope(Alyamani & Lemine, 2012).

#### Field Emission Transmission Electron Microscopy (FETEM)

In order to prepare the sample for FETEM examination, a drop of the sample was placed on carbon-coated copper grids, and then it was allowed to air dry overnight. Micrographs were obtained by using JOEL, Model: 2100 transmission electron microscope. This analytical technique is used to investigate the size, shape, structures and their electronic properties(Smith, 2015).

#### Fourier Transform Infrared [FTIR] spectroscopy

The FTIR spectrum of synthesized nanoparticles was analyzed using IR Affinity-1S instrumentIt is used to determine the nature of the functional groups that are associated with nanoparticles in biological extracts and the structural properties of the biological extract (Baudot et al., 2010).

#### MTT based cell viability assay against human breast cancer cell line MCF-7

The MTT Assay is a colorimetric assay to evaluate cell metabolic activity. Human breast cancer cell lines (MCF7) were maintained in DMEM with 10% FBS. To assess the cytotoxic potential of MaAgNP, 5×10^3^ cells were seeded in a 96-well microtiter plate (Thermo Nunc) with fresh 10% serum medium and incubated for 24 hours for their adherence. After adherence cells were treated with increasing concentrations of the MaAgNPs and AgNPs and allowed to grow for 48 hours at 37°C in a CO_2_ incubator maintaining 5% CO_2_ level. After 48 hours of incubation, 5 μl of MTT reagent (from 5 mg/ml stock) was added to each well in dark conditions, and the plate was incubated for 2 hours at 37°C under same condition as mentioned previously. The medium was then withdrawn, and 200 L of DMSO was added to dissolve the generated formazan crystal complex. Using the Tecan Microplate Reader Infinite 200 Pro, the intensity of the dissolved formazan crystals (purple colour) was measured at 570 nm with reference at 630 nm..

The IC_50_ value was calculated using GraphPad Prism software with the normalized absorbance percentage vs log concentration (Ganot et al., 2013).

#### Reactive oxygen species (ROS) detection assay

Utilizing 2’, 7’ dichlorodihydrofluorescein diacetate (DCF-DA) staining allowed for the monitoring of the production of hydrogen peroxide and superoxide radicals. DCF-DA is colourless and nonfluorescent unless it is oxidised by reactive oxygen species found within the cell.. 4 x 10^5^ cells were seeded for 24 hours and then treated with MaAgNP at IC_50_ concentration and incubated for 48 hours at 37°C in a CO_2_ incubator maintaining 5% CO_2_ level. After incubation cells were washed with 1X PBS and harvested with trypsin. Cells were finally resuspended in 1 ml PBS and stained with 10 μM DCF-DA. Stained cells were then incubated at 37°C for 45 mins. After incubation cells were washed again with PBS and resuspended in 1ml PBS and analyzed using a BD FACScalibur flow cytometer(Figueroa et al., 2018).

#### Inverted fluorescence microscopy analysis

For the fluorescence microscopic analysis, 4 x 10^5^ cells were seeded for 24 hours and then treated with MaAgNP at IC_50_ concentration and incubated for 48 hours at 37°C in a CO_2_ incubator maintaining 5% CO_2_ level. After incubation cells were stained with DCF-DA and analyzed under an inverted fluorescence microscope (Aleksandra Wojtala et al., 2014.).

### Statistical analysis

All data were expressed as mean ± SD of the experiment. Data analysis was conducted using Graphpad prism software. The data were analysed using a one-way analysis of variance (ANOVA) test in order to make comparisons for the MTT experiment. When the p-value was less than 0.001, the differences were considered to be statistically significant(Swift, 1997.).

## Results and Discussion

### Chemical synthesis of silver nanoparticles

The appearance of brown colour on the solution (Fig 1) which contain 1mM silver nitrate (AgNO_3_) indicated the synthesis of silver nanoparticles in the solution. Approximately 2mg of AgNPs were produced from 100 ml of 1mM AgNO_3_ solution by chemical synthesis.

**Fig 1:**
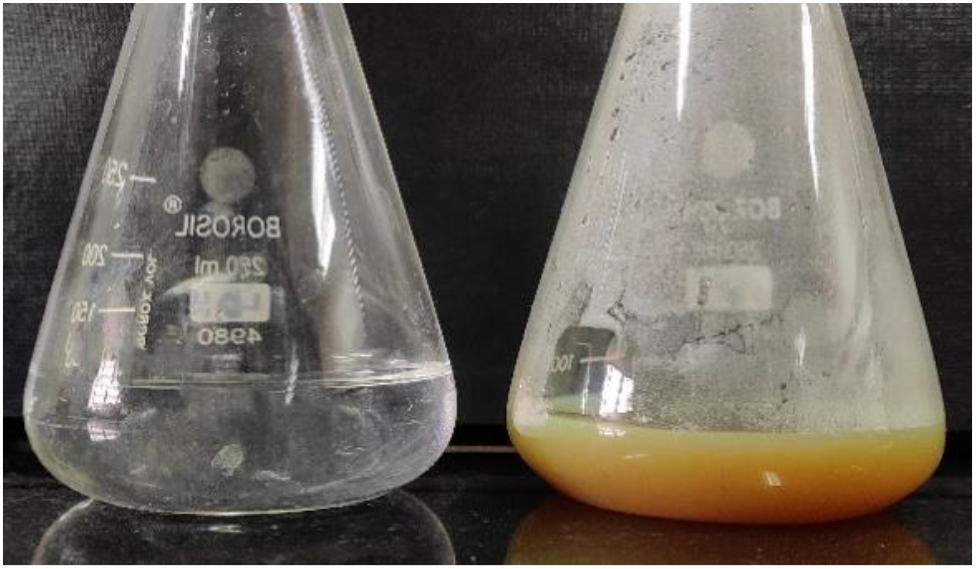
Visual observation of colour change in control and chemically synthesized silver nanoparticles.

### Mycosynthesis of silver nanoparticles

The reduction process results in the formation of AgNP, which may be visually noticed as a change in the colour of the solution from colourless to dark brown. This change serves as an indicator of the formation of AgNP (Fig 2). *Metarhizium anisopliae* filtrate exposed to silver nitrate exhibits dark brown colour used as a test. The dark brown colour can be because of the surface plasmon resonance effect and also due to the reduction of AgNO_3_. *M. anisopliae* filtrate without the exposure of silver nitrate was kept in control showed no changes in colour. After separation and purification, approximately 30 mg of silver nanoparticles were produced from 100ml of fungal filtrate containing 1mM AgNO_3_ that was a 15 times increase in the nanoparticles yield as compared to the chemical synthesis.

**Fig 2:**
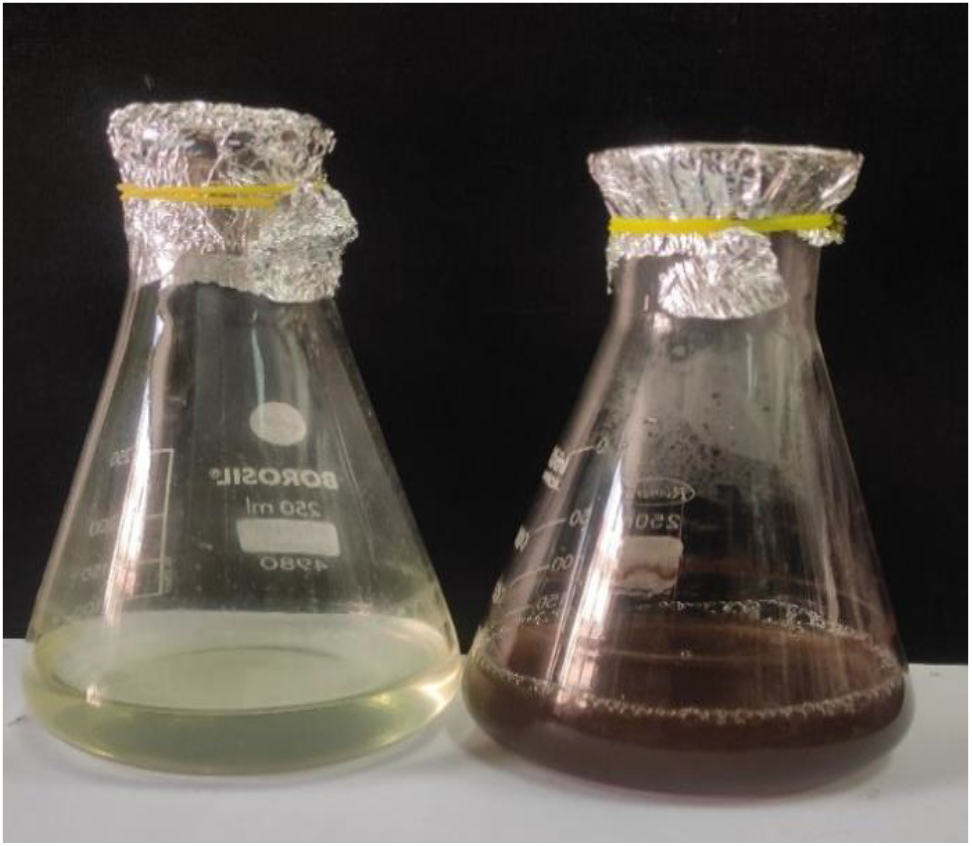
Visual observation of colour change in control and mycosynthesized silver nanoparticles.

### UV-VIS Spectroscopy analysis of silver nanoparticles

Nanoparticles of silver have their own distinct optical properties, which allow them to have a powerful interaction with certain wavelengths of light. The conduction band and the valence band of silver nanoparticles lay relatively near to one another, allowing electrons to travel freely between the two bands. Because of the mutual oscillations that occur between the electrons of nanoparticles when they are in resonance with a light wave, these free electrons have the potential to produce an absorption band known as surface plasmon resonance, or SPR (Maribel G. Guzmán et al., 2009). The UV-visible absorption spectra of the mycosynthesized silver nanoparticles (MaAgNP) and chemically synthesized (AgNP) were recorded (Fig. 3) both showing an absorption peak at around 420 nm.

**Fig 3:**
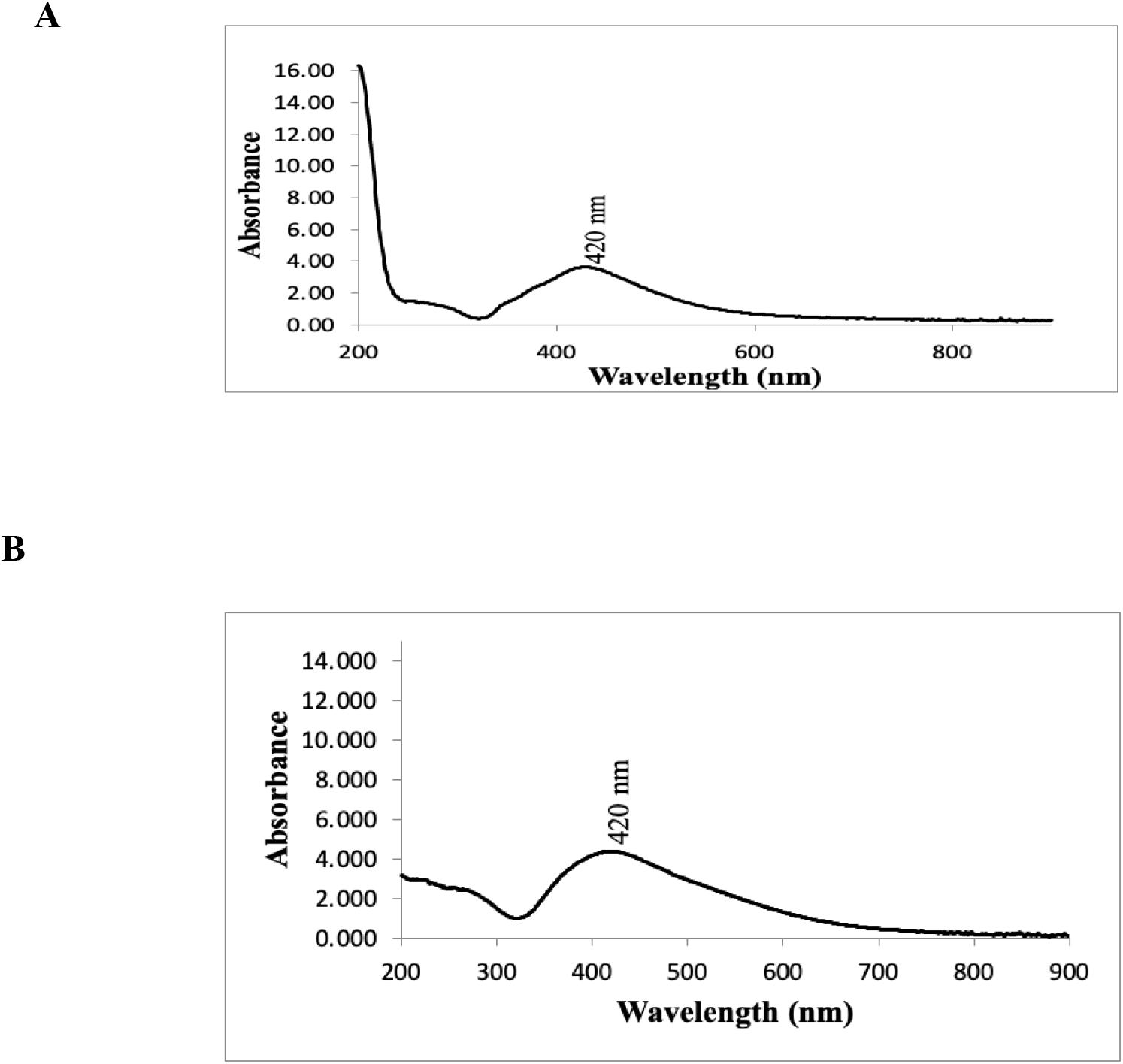
UV-Vis spectra of (A) chemically synthesized AgNP. (B) mycosynthesized silver nanoparticles.

### Field Emission Scanning Electron Microscopic analysis of MaAgNP

FESEM was employed to record the micrograph of mycosynthesized silver nanoparticles (MaAgNP). Scanning electron micrograph showed (Fig 4) the presence of aggregated rod-shaped uniformly distributed silver nanoparticles synthesized using *Metarhizium anisopliae*.

**Fig 4:**
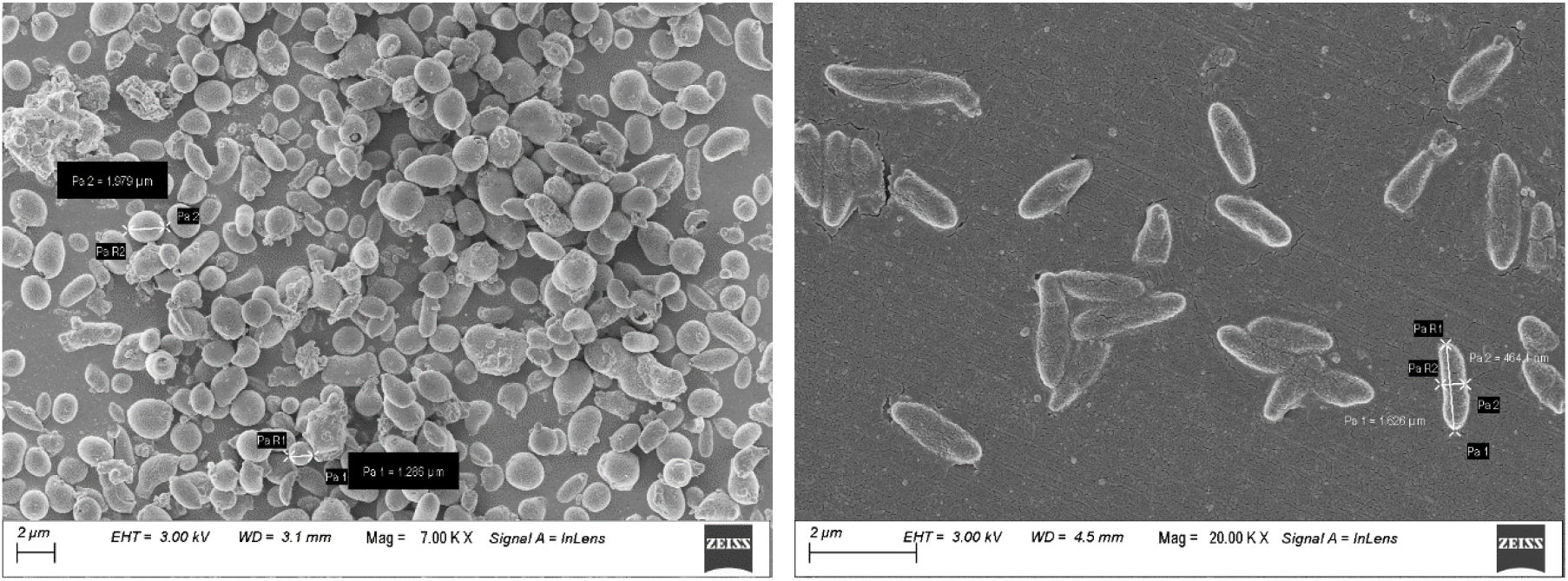
FESEM images of MaAgNP as powder sample (left) and liquid sample (right).

### Field Emission Transmission Electron microscopic analysis of MaAgNP

The silver nanoparticles produced from *Metarhizium anisopliae* were characterized using Field Emission Transmission Electron Microscope (FETEM). The images obtained from FETEM (Fig. 5) showed the nanoparticles having a spherical shaped structure and a size range of 5-20 nm. Nanoparticle aggregate are rod shape and have ~ 2.18 μm size (Fig. 5), which can be compared to the FESEM images in Fig 4. The selected area electron diffraction (SAED) pattern (Fig. 5) shows that the nanoparticles obtained possess polycrystallinity.

**Fig 5:**
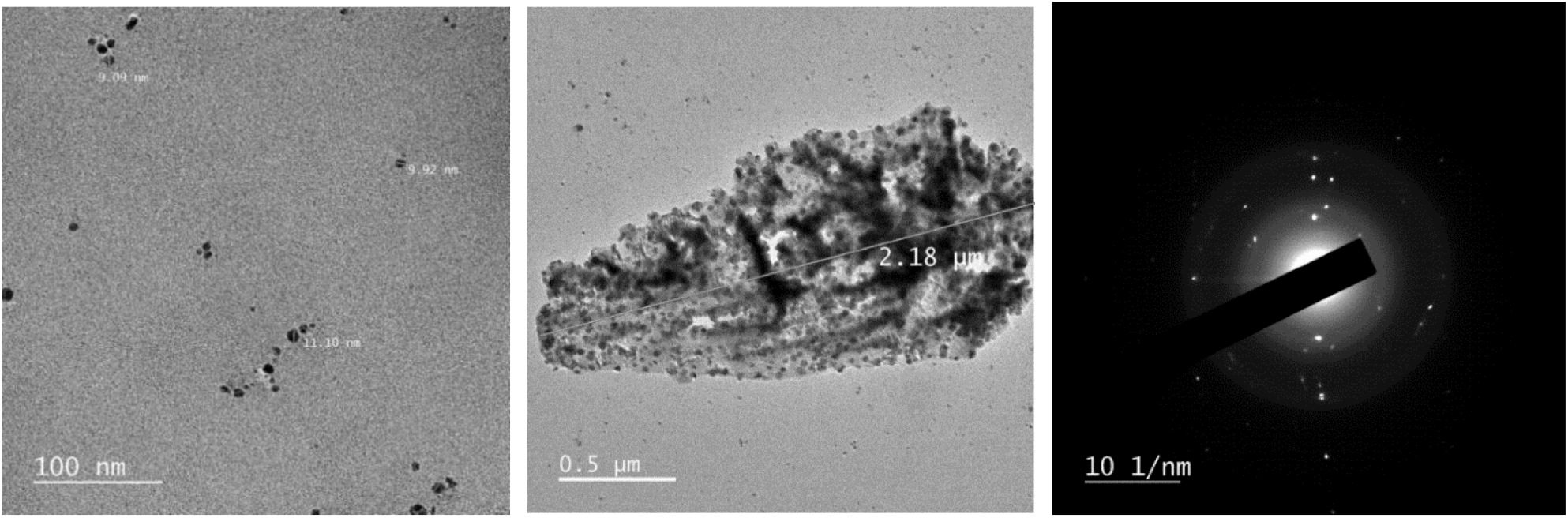
FETEM images of MaAgNP as individual nanoparticle (left) AgNP aggregates (middle) and SAED pattern of a single nanoparticle (right).

### Fourier Transform Infrared Spectroscopy (FTIR) analysis of MaAgNP

The samples of silver nanoparticles were examined using FTIR in order to determine the probable biomolecules that were responsible for the reduction of Ag+ ions to AgNPs induced by the fungal filtrate.. In MaAgNP the significant peaks were located mainly at 3246, 1539, 952, and 540 cm^-1^ corresponding to alcohol (-OH), nitro compound (N-O), alkene (C=C), and halo compound (C-I) (Fig. 6).

**Fig 6:**
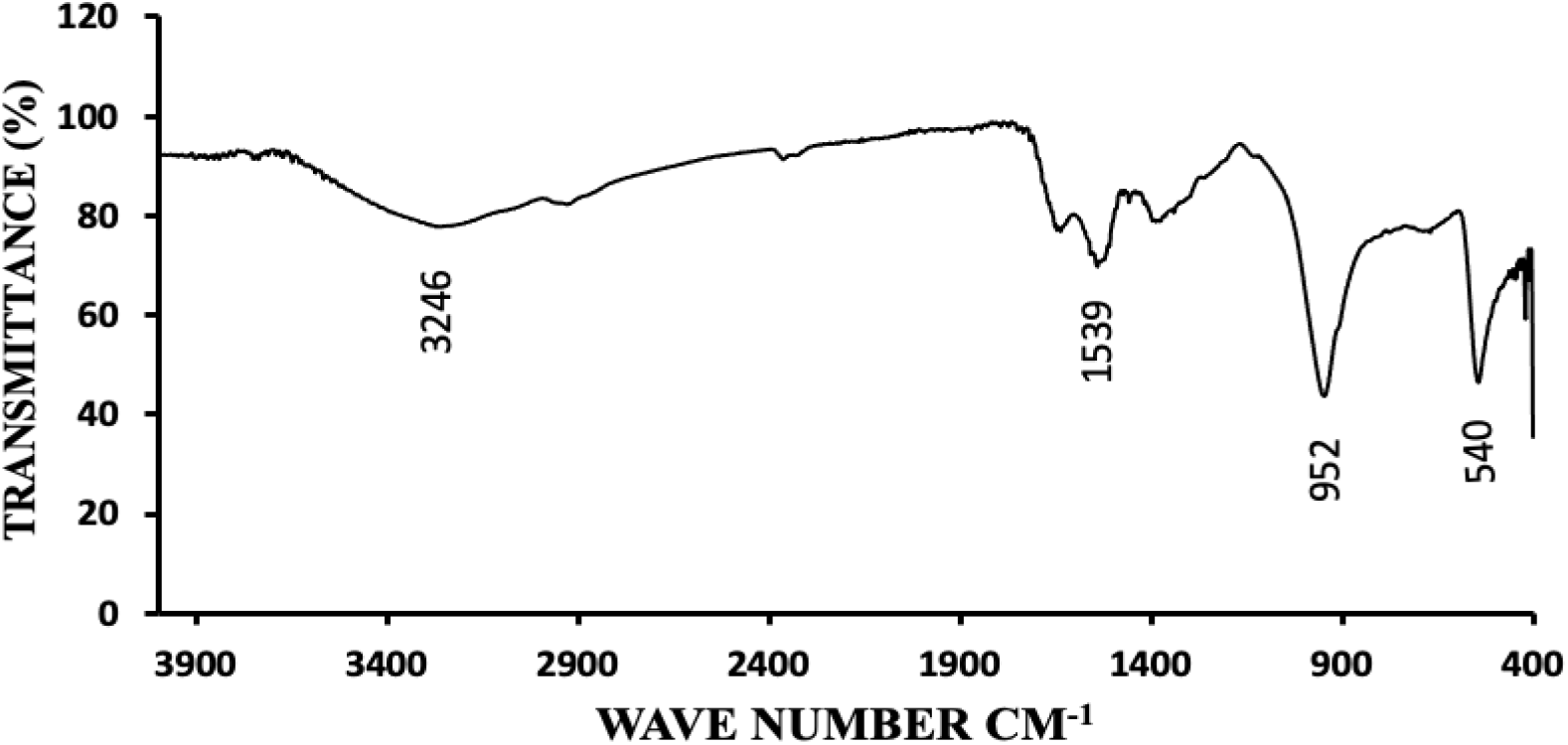
FTIR analysis of MaAgNP.

### MTT based cell viability assay against human breast cancer cell line MCF-7

The MTT assay was used to assess whether chemically produced silver nanoparticles and myco-synthesized silver nanoparticles display any cytotoxicity on MCF-7 cell lines by measuring the cellular activity as a sign of cell damage or cytotoxicity. This was done using the MTT method. It is a calorimetric assay, and it is based on the capability of NADPH-dependent cellular oxidoreductase enzymes to convert yellow tetrazolium dye (MTT) to its insoluble formazan crystal, which is purple in colour. So, if the cells are metabolically active then they will be more purple compared to dying cells or dead cells. In this assay, MaAgNPs showed gradual decrease in the cell viability with increasing concentration of the nanoparticles dosage. The half maximal inhibitory concentration (IC_50_) value of MaAgNP for MCF7 cell lines was found to be 16.50μg/ml. Whereas, the chemically synthesized silver nanoparticles (AgNPs) showed a constatnt viable cell population with increasing nanoparticles concentration up to 100 μg/ml and no IC_50_ value could be determined as it could not reach the half maximal population inhibition even with the highest concentration used here (Fig.7).

**Fig 7:**
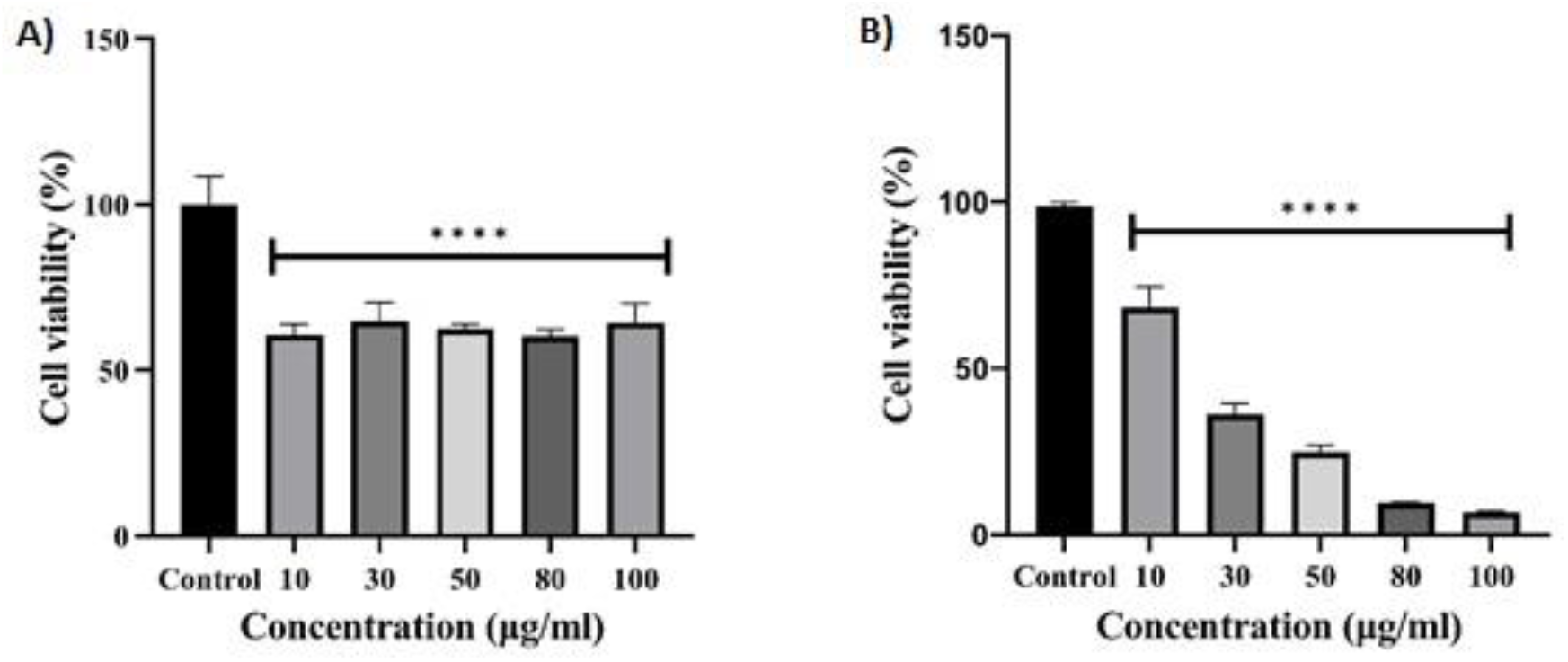
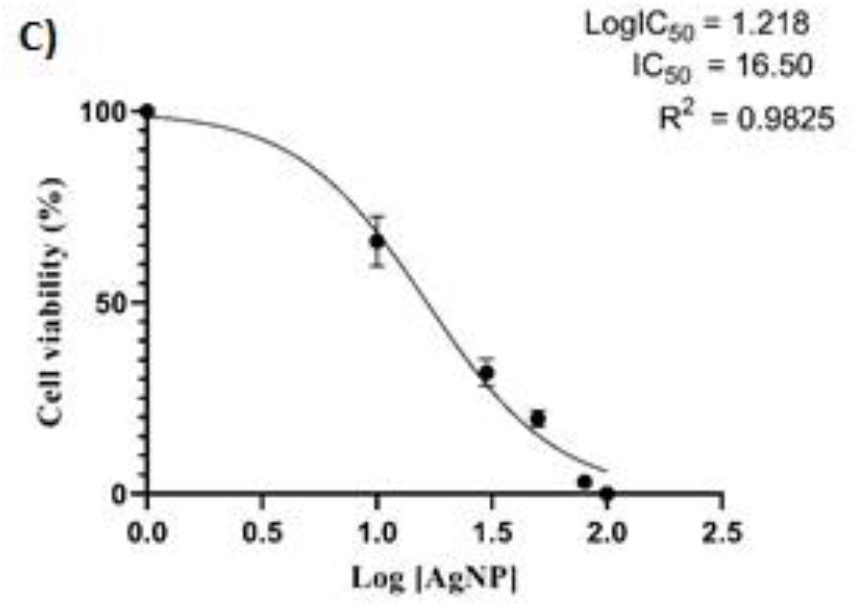
MTT cell viability assay showing the cell viability (%) (A) % survival of cells after the treatment of chemically synthesized AgNP, (B) % survival of cells after the treatment of MaAgNP (C) Calculation of IC_50_ value for MaAgNP treated cells.

### Reactive oxygen species (ROS) detection assay

2’, 7’-dichlorodihydrofluorescein diacetate (H2DCF-DA) fluorescent probe is used to detect the reactive oxygen species produced in the cell. Upon oxidation by ROS, the nonfluorescent H2DCFDA is converted to the highly fluorescent 2’, 7’-dichlorofluorescein (DCF). Only dead or dying cells produce reactive oxygen species. Here fluorescence microscopy is employed to assess probe fluorescence emission. From the images (Fig. 8) it is evident that untreated cells were able to maintain the rigid morphological structure as observed in the bright field and do not produce any reactive oxygen species, hence producing no colour under fluorescence microscopy. The treated cells showed an absence of adherence and altered cellular morphology. The reactive oxygen species produced by the cells were oxidized the dye and produced green fluorescent in the treated cells. Further the flow cytometric analysis results (Fig. 9) showed that there is a 0.5-fold increment in ROS generation in the treated samples compared to the untreated one which is the indication of cellular apoptosis (Simon et al., 2000).

**Fig 8:**
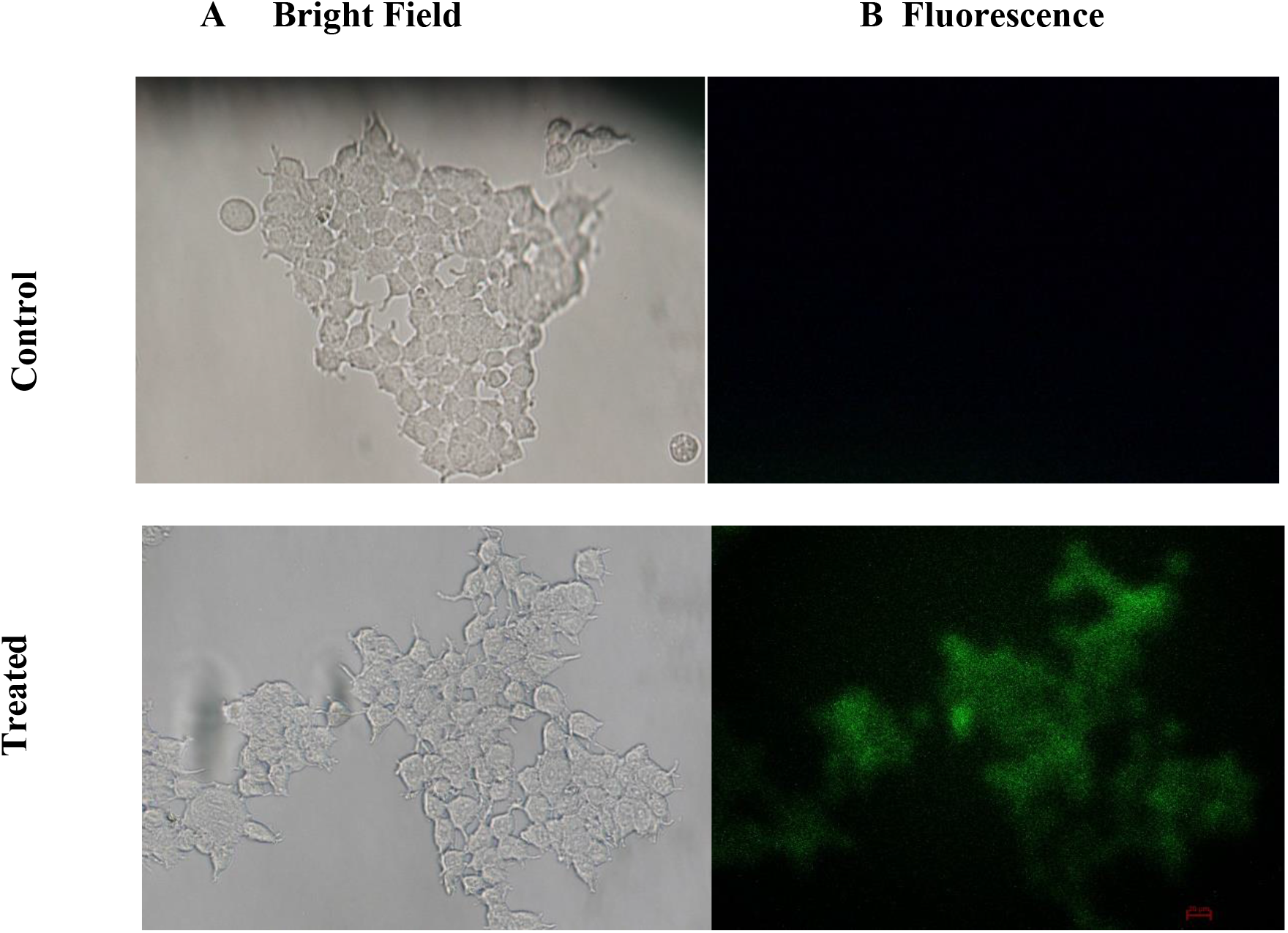
Fluorescence microscopy analysis of ROS generation in MaAgNPs treated MCF7 cells cmpared to the untreated cells (negative control).

**Fig 9:**
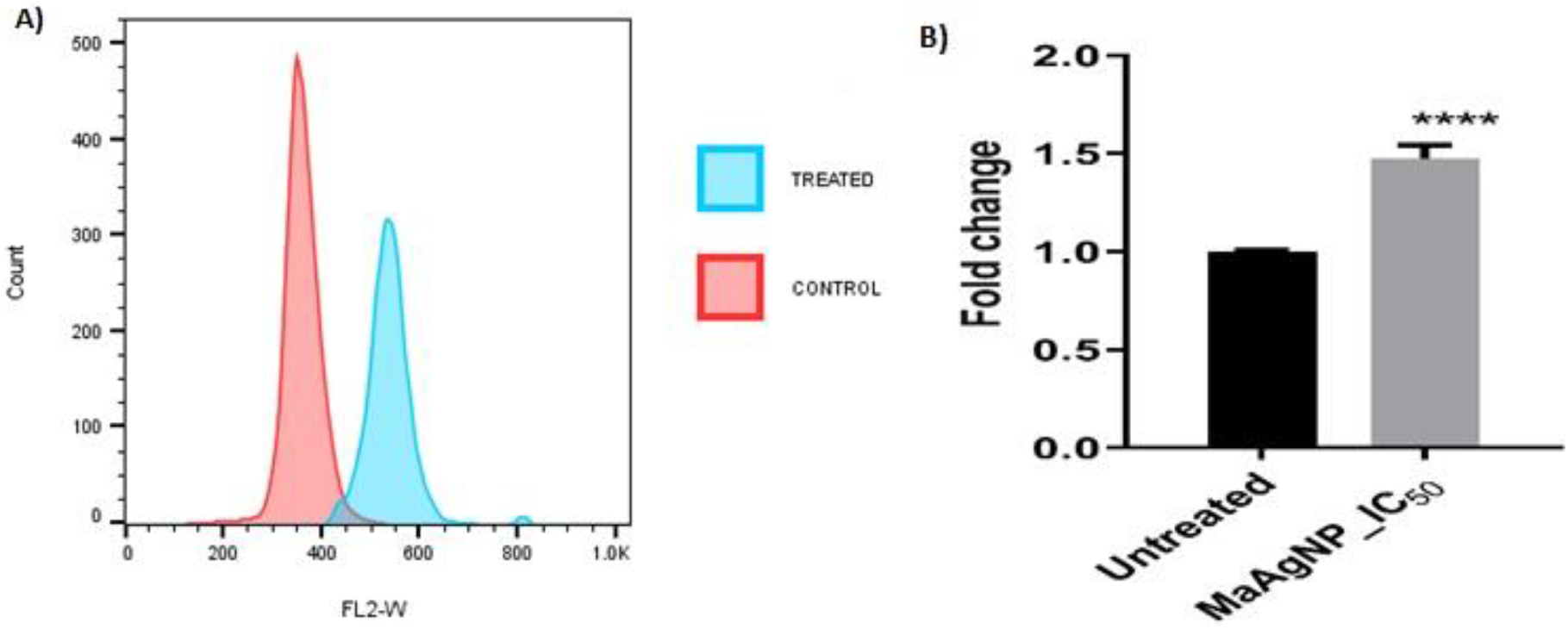
(A) Flow cytometric analysis of ROS generation in MaAgNPs treated MCF7 cells compared to the untreated cells (B) Quantitative analysis of ROS generation.

## Conclusions

Studies related to the usage of silver nanoparticles are becoming multifold day by day because of their multifaceted potential for the treatment of various diseases or drug delivery and controlling vector of disease-causing agents. Studies related to AgNP showed that it can induce cytotoxicity in microbial pathogens as well as cancer cells ((Burduşel et al., 2018). In this study, silver nanoparticles were successfully synthesized using *M. anisopliae*. The mycosynthesis method gives a high yield compared to chemical synthesis and also it does not produce any hazardous by-products and no need to use costly toxic chemicals for synthesis. The characterization of nanoparticles showed that the average size of a single nanoparticle is around 20 nm which is ideal to be delivered in cells. The analysis of nanoparticle cytotoxicity on cancer cell lines was analysed by cell viability assay on the cell line, and it revealed that mycosynthesized silver nanoparticles showed a prominent effect on viability when compared with the chemically synthesized silver nanoparticles. Further ROS assay also proved that the MaAgNP treated cells produced reactive oxygen species which indicates activation of cell death via apoptosis.

This study clearly details the significant effect of mycosynthesized silver nanoparticles on cancer cell lines, which could be exploited further for the development of potential cancer drugs based on fungus and fungal secondary metabolites.

## Abbreviations

AgNP: silver nanoparticle
FESEM: Field emission scanning electron microscope
FETEM: Field emission transmission electron microscope
FTIR: Fourier transform infrared spectroscopy
MaAgNP: *Metarhizium anisopliae* silver nanoparticle
MCF-7: Michigan cancer foundation −7
ROS: Reactive oxygen species

## Acknowledgements

This research was supported by Department of Biosciences and Bioengineering, IIT Guwahati.

## Statements and Declarations

### Conflict of interest

The authors declare that they have no known competing financial interests or personal relationships that could have appeared to influence the work reported in this paper.

### Author contribution

Design of the work: Sujisha S Nambiar

Data collection: Sujisha S Nambiar, Arman Mohanty, Arupam Patra

Data analysis: Sujisha S Nambiar, Arman Mohanty, Arupam Patra

Data interpretation: Sujisha S Nambiar, Arman Mohanty, Arupam Patra

Manuscript writing: Sujisha S Nambiar

Supervision: Gurvinder Kaur Saini

## Notes

### Competing Interest Statement

The authors have declared no competing interest.

## REFERENCES

Abou El-Nour, K. M. M., Eftaiha, A., Al-Warthan, A., & Ammar, R. A. A. (2010). Synthesis and applications of silver nanoparticles. In Arabian Journal of Chemistry (Vol. 3, Issue 3, pp. 135–140). https://doi.org/10.1016/j.arabjc.2010.04.008

Aleksandra Wojtala, Massimo Bonora, Dominika Malinska, Paolo Pinton, Jerzy Duszynski, & Mariusz R. Wieckowski. (n.d.). Chapter Thirteen - Methods to Monitor ROS Production by Fluorescence Microscopy and Fluorometry.

Alyamani, A., & Lemine, O. M. (n.d.). FE-SEM Characterization of Some Nanomaterial. www.intechopen.com

Amerasan, D., Nataraj, T., Murugan, K., Panneerselvam, C., Madhiyazhagan, P., Nicoletti, M., & Benelli, G. (2016). Myco-synthesis of silver nanoparticles using Metarhizium anisopliae against the rural malaria vector Anopheles culicifacies Giles (Diptera: Culicidae). Journal of Pest Science, 89(1),249–256. https://doi.org/10.1007/s10340-015-0675-x

Baudot, C., Tan, C. M., & Kong, J. C. (2010). FTIR spectroscopy as a tool for nano-material characterization. Infrared Physics and Technology, 53(6), 434–438. https://doi.org/10.1016/j.infrared.2010.09.002

Burduşel, A. C., Gherasim, O., Grumezescu, A. M., Mogoantă, L., Ficai, A., & Andronescu, E. (2018). Biomedical applications of silver nanoparticles: An up-to-date overview. In Nanomaterials (Vol. 8, Issue 9). MDPI AG. https://doi.org/10.3390/nano8090681

Chandra, J. H., Raj, L. F. A. A., Namasivayam, S. K. R., & Bharani, R. S. A. (2013). Improved pesticidal activity of fungal metabolite from nomureae rileyi with chitosan nanoparticles. Proceedings of the International Conference on “Advanced Nanomaterials and Emerging Engineering Technologies”, ICANMEET 2013, 387–390. https://doi.org/10.1109/ICANMEET.2013.6609326

David L Hawksworth, & Robert Lucking. (n.d.). Fungal Diversity Revisited: 2.2 to 3.8 Million Species.

Feofilova, E. P. (2001). The Kingdom Fungi: Heterogeneity of Physiological and Biochemical Properties and Relationships with Plants, Animals, and Prokaryotes (Review). Applied Biochemistry and Microbiology 37, 124–137.

Figueroa, D., Asaduzzaman, M., & Young, F. (2018). Real time monitoring and quantification of reactive oxygen species in breast cancer cell line MCF-7 by 2’,7’–dichlorofluorescin diacetate (DCFDA) assay. Journal of Pharmacological and Toxicological Methods, 94, 26–33. https://doi.org/10.1016/j.vascn.2018.03.007

Ganot, N., Meker, S., Reytman, L., Tzubery, A., & Tshuva, E. Y. (n.d.). Anticancer Metal Complexes: Synthesis and Cytotoxicity Evaluation by the MTT Assay.

Gudikandula, K., & Charya Maringanti, S. (2016). Synthesis of silver nanoparticles by chemical and biological methods and their antimicrobial properties. Journal of Experimental Nanoscience, 11(9),714–721. https://doi.org/10.1080/17458080.2016.1139196

Maribel G. Guzmán, Jean Dille, & Stephan Godet. (2009). Synthesis of silver nanoparticles by chemical reduction method and their antibacterial activity. International Journal of Chemical and Biomolecular Engineering 2:3.

Sastry, M., Ahmad, A., Khan Islam Maksudul, & Kumar, R. (n.d.). Biosynthesis of metal nanoparticles using fungi and actinomycete. https://www.researchgate.net/publication/228550063

Shameli, K., Ahmad, M. bin, Yunus, W. M. Z. W., Ibrahim, N. A., Gharayebi, Y., & Sedaghat, S. (2010). Synthesis of silver/montmorillonite nanocomposites using γ-irradiation. International Journal of Nanomedicine, 5(1), 1067–1077. https://doi.org/10.2147/IJN.S15033

Simon, H.-U., Haj-Yehia, A., & Levi-Schaffer, F. (2000). Role of reactive oxygen species (ROS) in apoptosis induction. In Apoptosis (Vol. 5). Kluwer Academic Publishers.

Smith, D. J. (2015). CHAPTER 1: Characterization of nanomaterials using transmission electron microscopy. In RSC Nanoscience and Nanotechnology (Vols. 2015-January, Issue 37, pp. 1–29). Royal Society of Chemistry. https://doi.org/10.1039/9781782621867-00001

Swift, M. L. (n.d.). GraphPad Prism, Data Analysis, and Scientific Graphing.http://www.graphpad.com

Verena seidl. (n.d.). Chitinases of filamentous fungi: a large group of diverse proteins with multiple physiological functions.

Zhang, X. F., Liu, Z. G., Shen, W., & Gurunathan, S. (2016). Silver nanoparticles: Synthesis, characterization, properties, applications, and therapeutic approaches. In International Journal of Molecular Sciences (Vol. 17, Issue 9). MDPI AG. https://doi.org/10.3390/ijms17091534

